# Biomechanical Evaluation of Temporomandibular Joint Implants and Periprosthetic Bone under Unilateral and Bilateral Clenching

**DOI:** 10.1101/2024.08.26.609607

**Authors:** Rajdeep Ghosh, Girish Chandra, Vivek Verma, Kamalpreet Kaur, Ajoy Roychoudhury, Sudipto Mukherjee, Anoop Chawla, Kaushik Mukherjee

## Abstract

To ensure the long-term success of temporomandibular joint (TMJ) implants, it is imperative to understand their biomechanical performances. This study aims to compare the biomechanical performance of two stock implants (narrow and standard) under unilateral and bilateral clenching during both osseointegrated and non-osseointegrated conditions. Finite element models of a human mandible were developed from QCT data, with the left TMJ being replaced by the implants. Six clenching tasks were simulated to evaluate stress and strain distributions in the mandible and implants. Ipsilateral clenching produced higher mandibular strains, while contralateral clenching generated larger implant stresses. Furthermore, intercuspal biting was found to have produced the highest strain (1750-1880 µε) and stress (∼17 MPa) in the mandible. Osseointegration reduced stresses (up to 0.14 MPa) and strains (up to 30 µε) in mandible as well as stresses in mandibular components (up to 48 MPa) and screws (up to 71 MPa). However, during non-osseointegrated conditions, stresses in cortical bone were higher for standard TMJ implant as compared to narrow implant. This suggests possible preference of narrow implant over standard ones.

## 1. Introduction

A wide range of disorders, primarily intra-articular pathologies, and trauma ^1–3^, can affect the temporomandibular joint (TMJ), causing clinical symptoms such as pain, restricted joint movement, malocclusion, and functional difficulty during deglutition and speech ^4,5^. TMJ replacement with an alloplastic joint may be required for patients who do not respond well to conservative and non-surgical treatment ^1^. Despite the short-term successes associated with the use of TMJ implants, the long-term implications of commercially available implants are still not clear to the scientific community and clinicians ^6–9^.

The patient-specific TMJ Concepts system (Stryker, Ventura, CA, USA) and the stock Zimmer Biomet Microfixation systems (Jacksonville, FL, USA) are the two primary US-FDA-approved TMJ replacement systems now available globally^1^. Stock implants are widely used due to their affordability and shorter delivery times, making them the preferred choice in clinical practice^10^. Biomet Microfixation system^®^ designed stock mandibular prosthesis comes in two shapes: standard (45 mm, 50 mm and 55 mm) and narrow (45 mm and 50mm) with varying sizes depending on the length of the mandibular body. The choice between standard and narrow implants depends on factors such as patient anatomy, surgical requirements, and the extent of joint reconstruction needed. A report from a recent meta-analysis by Zou et al.^11^ revealed comparable outcomes between stock and patient-specific prostheses in terms of reduced pain score and improvement in function score, diet score, and maximum incisal opening. However, these implants (especially stock implants) are vulnerable to long-term failure risks primarily owing to implant design, poor material selection, improper surgical techniques etc.^7,8^. Therefore, further research is needed to achieve a longer life span for existing TMJ implants ^6,11^.

The human mandible experiences different physiological loads throughout its lifetime. Although there have been several *in silico* studies on biomechanical assessment of healthy ^12–18^ and reconstructed human mandibles ^19–27^, only a few studies have investigated reconstructed human mandibles with TMJ implants ^22,24,25^. Notable among them is the study by Pinheiro et al. ^24^ which reported the strain and deformation across the human mandible implanted with a patient-specific TMJ implant under three clenching tasks. Their study considered a frictionless interface between the bone-implant and condyle-fibrocartilage. A frictionless interface typically indicates an extremely smooth interface which offers no resistance against relative movement. While this might be true for the condyle-fibrocartilage joint, the bone-implant interfaces are typically modelled as frictional (non-osseointegrated) ^28–30^ or bonded (osseointegrated) ^29,31,32^. Additionally, such simplified bone-implant interface modelling would also influence the stress-strain behaviour across the mandible and TMJ implant.

To the authors’ knowledge, there is hardly any study investigating the influence of different stock TMJ implants on load transfer across the human mandible under both unilateral and bilateral clenching activities. The present study is thus aimed at gaining an insight into the biomechanical performance of two stock TMJ implants (standard and narrow types by Zimmer Biomet Microfixation systems^®^) under unilateral and bilateral clenching tasks, representing typical loading scenarios encountered during mastication. While focusing on controlled biomechanical evaluation, this study also investigates the influence of bone-implant interface conditions, representing osseointegrated and non-osseointegrated states, to simulate variations commonly observed in clinical outcomes.

## 2. Materials and Methods

Quantitative computed tomography (QCT)-dataset (512 x 512 pixels; voxel size of 0.439 × 0.439 × 0.75 mm^3^) of an anonymized 20-year-old healthy male subject was obtained to develop the model of the intact mandible. A 3-D surface model of the intact mandible was reconstructed in Materialise Mimics v25.0 (Materialise Inc., Leuven, Belgium). Finite element (FE) mesh was generated in Hypermesh v2022 (Altair Engineering Inc., Troy, MI, USA), whereas FE simulation and further post-processing were performed in Ansys APDL v2022 R2 (ANSYS Inc., Canonsburg, PA, USA).

### 2.1 Development of Intact FE model

The human mandible (Figure 1) has the following major components: cortical, cancellous, periodontal ligament (PDL), teeth, articular fibrocartilage, and temporal bone (represented by cubic block). Due to considerable influence of PDL layer on stress-strain distribution in human mandible ^14^, it was decided to model the PDL (∼ 0.3 mm thickness^15^) and this was achieved by offsetting the interface between the cancellous bone and the teeth. PDL was further divided into three subcomponents along the root length: coronal, middle, and apical thirds due to their region-specific distinct mechanical properties^12,19,26^. All components were meshed with 10-noded tetrahedral elements (with edge lengths from 0.3 mm to 2 mm) by fulfilling appropriate mesh quality parameters such as aspect ratio (<5), Jacobian ratio (>1), etc. A mesh convergence study was performed with an ascending number of elements from coarse (305,774 elements), to medium (3,403,988 elements), to fine (7,054,373 elements) element sizes on the intact model. However, difference in maximum principal stress in the mandible was predicted to be 5% between coarse and medium meshed models, while 0.4% between the medium and fine meshes. Therefore, the medium meshed model with element sizes ranging between 0.5 mm to 1.5 mm (3,403,988 elements) was chosen as the optimum FE model for further investigation in this study.

**Figure 1.**
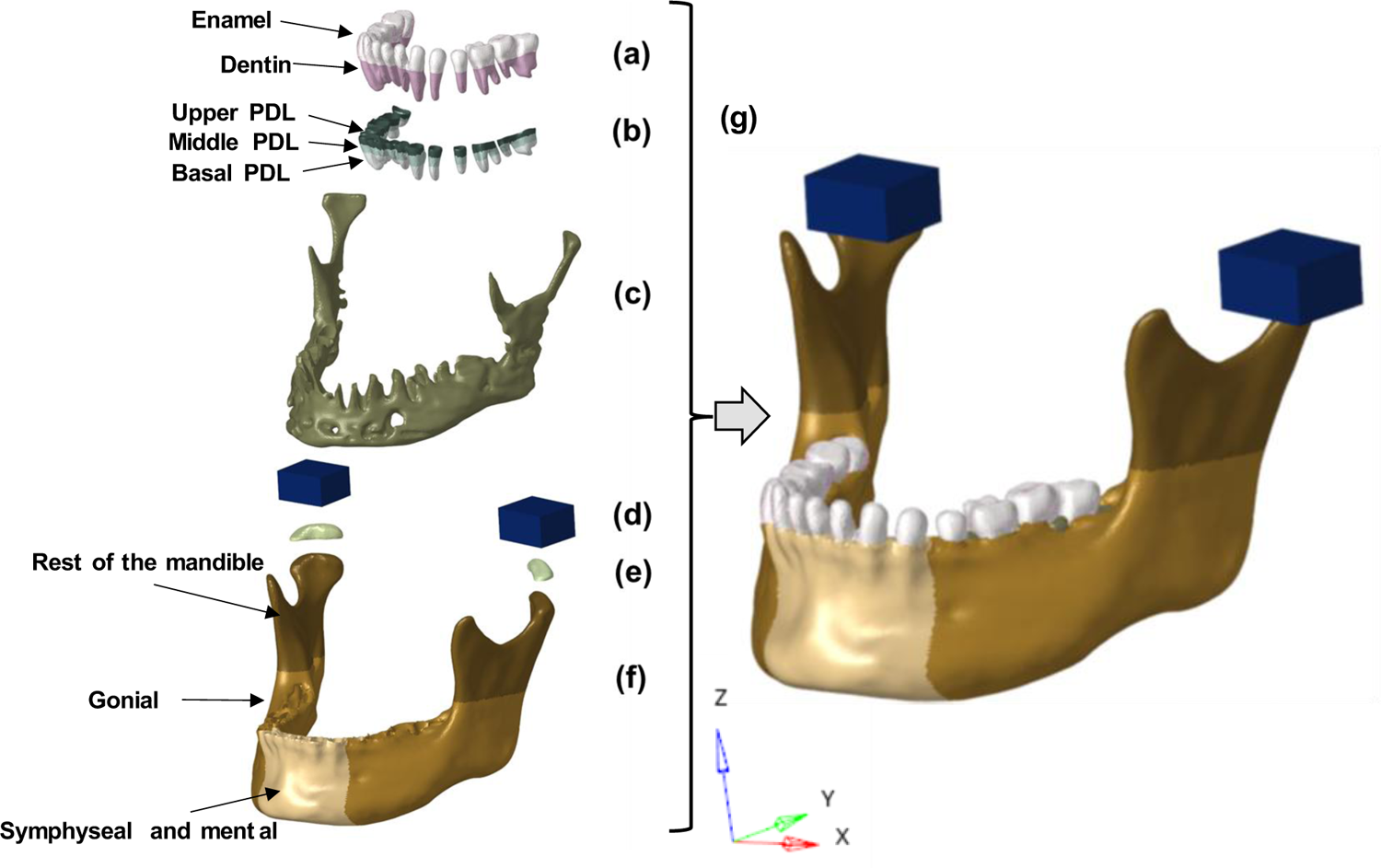
Different components of the developed mandible model: (a) teeth with enamel and dentin; (b) periodontal ligament (PDL) comprising of upper, middle, and basal PDL; (c) cancellous bone; (d) blocks representing temporal bone; (e) articular fibrocartilage; (f) cortical bone with three major regions: symphyseal and mental, gonial, rest of the mandible; and (g) the complete mandible model.

### 2.2 Development of Implanted FE models

The 3D models of the mandibular components of TMJ implants (Figure 2 - a & b) were developed from the micro-CT images (Xtreme CT II, Scanco Medical AG, Switzerland, with scanning parameters: 491 x 256 pixels; isometric voxel size of 60 μm) of narrow and standard stock implants of size 45 mm (Zimmer Biomet Microfixation system, Jacksonville, FL, USA). Based on Biomet mandibular screws ^33^, bi-cortical screws were modelled as simplified cylinders (Φ2.7 mm) with lengths ranging from 8 – 12 mm. The models of the intact mandible and TMJ implant were imported into Rhinoceros^®^ software (Rhinoceros V7.0, Robert McNeel & Associates, Seattle, USA) for osteotomy and virtual implantation based on clinical recommendations. Similarly to the intact models, a mesh convergence study was also performed with an ascending number of elements from coarse (narrow TMJ: 284,196 elements; standard TMJ: 296,378 elements), to medium (narrow TMJ: 2,950,186 elements; standard TMJ: 3,089,465 elements), to fine (narrow TMJ: 6,763,941 elements; standard TMJ: 6,876,264 elements) meshes on implanted models. Difference in maximum principal stresses in both the implanted mandibles were ∼ 7% between coarse and medium meshed models, and ∼ 0.5% between the medium and fine meshes. Therefore, the medium meshed models with element sizes ranging between 0.5 mm to 1.5 mm (narrow TMJ: 2,950,186 elements; standard TMJ: 3,089,465 elements) were chosen as the optimum FE models for further investigation in this study (Figure 2- c & d). For implanted model, the same temporal bone block represented the temporal bone resurfaced with fossa component.

**Figure 2.**
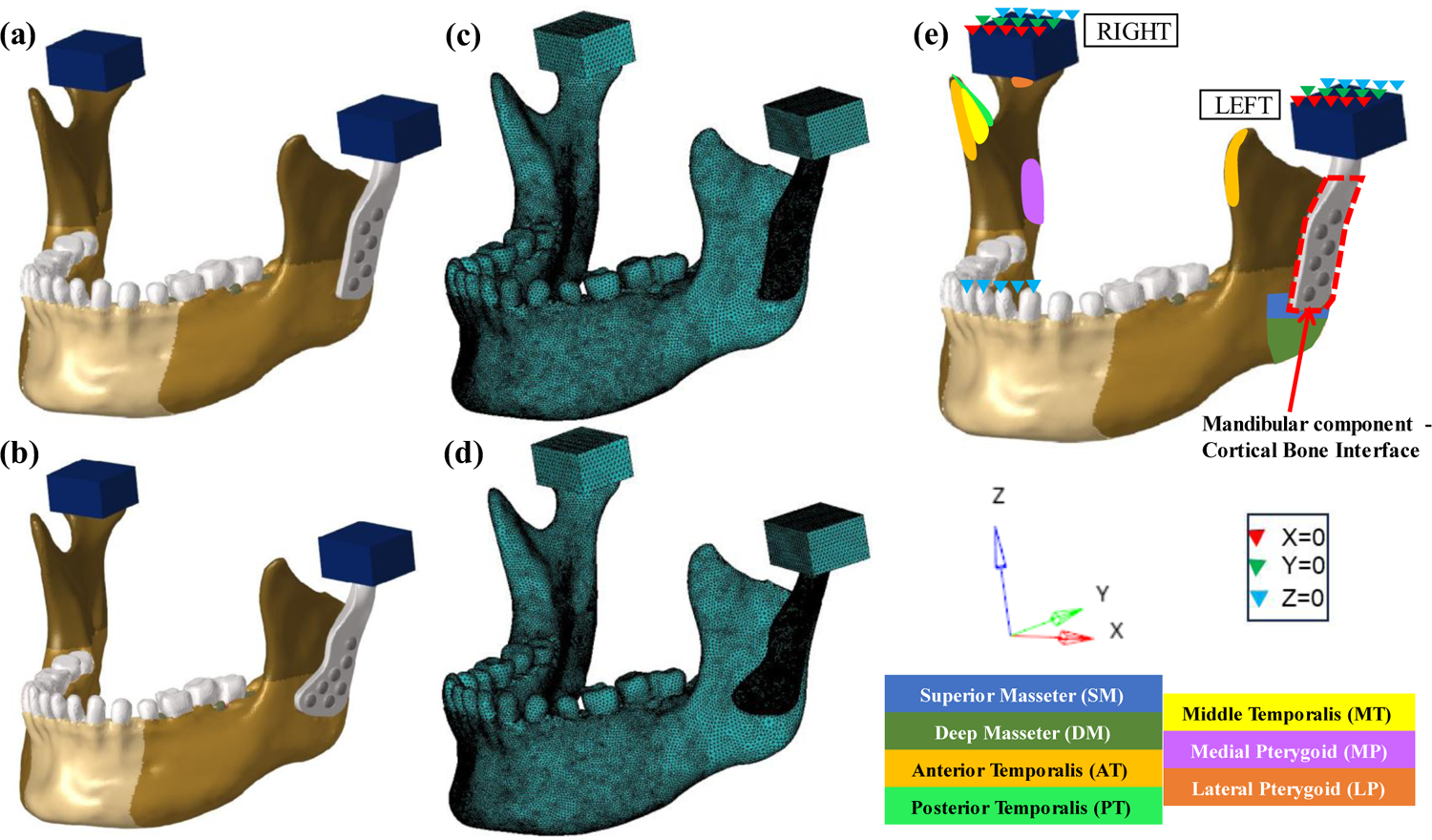
Developed FE models of implanted mandibles: (a, c) Narrow TMJ Implant (b, d) Standard TMJ Implant (e) muscle attachment sites and representative boundary conditions for incisor (INC) loading condition.

### 2.3 Material Properties

In this study, linear elastic orthotropic material properties were assigned to different regions of mandible. Cortical bone was modelled as a region-specific orthotropic material^12^ to better replicate the inhomogeneous and anisotropic nature of bone. Assigning region-specific orthotropic material properties to the cortical bone improves the model’s fidelity and ensures that stress-strain distributions are representative of reality^12^. In particular, the symphyseal and mental regions are known to exhibit differences in stiffness and strength compared to other regions of the mandible^12^. Cancellous bone, however, was modelled as homogeneous orthotropic material ^12^. All other components such as teeth, PDL, fibrocartilage and implant were modelled as linear elastic isotropic material ^19,26^. The temporal bone (in intact model) and resurfaced temporal bone with fossa component (in implanted model) were modelled as a rigid body ^19^ since the deformation in these components during mastication was out of the scope of the present study. Table 1 shows the material properties used in this study for different components. For the implanted mandible, the mandibular component and the screws were assumed to be made of medical grade Ti-6Al-4V extra-low interstitial (ELI).

**Table 1:** Material properties of the components of the FE model^12^.

| Components | Young's Modulus<br>(GPa) |  |  | Poisson's Ratio |  |  | Shear Modulus<br>(GPa) |  |  |
| --- | --- | --- | --- | --- | --- | --- | --- | --- | --- |
| | $E_x$ | $E_y$ | $E_z$ | $\nu_{xy}$ | $\nu_{yz}$ | $\nu_{xz}$ | $G_{xy}$ | $G_{yz}$ | $G_{xz}$ |
| **Cortical** |  |  |  |  |  |  |  |  |  |
| Symphyseal and mental | 23 | 10 | 15 | 0.3 | 0.3 | 0.3 | 4.8 | 3.6 | 6.2 |
| Gonial angle region | 20 | 11 | 12 | 0.3 | 0.3 | 0.3 | 4.8 | 5.3 | 6.0 |
| Rest of the cortical | 17 | 6.9 | 8.2 | 0.3 | 0.3 | 0.3 | 2.8 | 2.9 | 4.6 |
| **Cancellous** | 0.96 | 0.32 | 0.39 | 0.3 | 0.3 | 0.3 | 0.09 | 0.13 | 0.17 |
| **Enamel** | 80 | - | - | 0.3 | - | - | - | - | - |
| **Dentin** | 17.6 | - | - | 0.25 | - | - | - | - | - |
| **Periodontal ligament** |  |  |  |  |  |  |  |  |  |
| Coronal third | 0.0032 | - | - | 0.45 | - | - | - | - | - |
| Middle third | 0.0027 | - | - | 0.45 | - | - | - | - | - |
| Apical third | 0.0025 | - | - | 0.45 | - | - | - | - | - |
| **Fibrocartilage** | 0.006 | - | - | 0.47 | - | - | - | - | - |
| **Mandibular component and Screws**<br>**(Ti-6Al-4V ELI)** | 113.8 | - | - | 0.342 | - | - | - | - | - |

### 2.4 Loading and Boundary Conditions

The following six biting tasks (Table 2) were considered in this study: incisal (INC), intercuspal (ICP), right molar (RMOL), left molar (LMOL), right group (RGF), and left group (LGF) biting. Muscle activity of seven muscles (Figure 2-e) ^34^ namely, two masseter (SM/DM), two pterygoid (MP/LP), and three temporalis muscles (AT/MT/PT) were considered in this study. These muscle forces are also shown for better illustration in Figure 3 with representative boundary conditions and approximate loading direction and magnitude (proportional to its arrow length). However, the influence of lateral pterygoid was neglected on the implanted side of the implanted model as total joint replacement requires excision of condyle where the lateral pterygoid muscle is attached. In addition, the present study hypothesized that all the muscles reattached successfully post-implantation. The top surface of temporal bone blocks was fixed in all directions ^12,34^. The bite points corresponding to each biting tasks were constrained in Z-direction ^24^. To investigate the influence of bone-implant interface conditions on stress-strain distribution across the bone and implant, both osseointegrated (OI) and non-osseointegrated (NOI) bone-implant interface conditions have been considered here. To model the osseointegrated (OI) condition, the interface between mandibular component and cortical bone was bonded whereas a frictional interface (µ = 0.3) between mandibular component and cortical bone was considered to mimic the non-osseointegrated (NOI) condition. Furthermore, the frictionless contact between articular fibrocartilage and mandibular condyle was defined in the intact and implanted mandibles. Frictional contact (µ = 0.3) was considered between the head of the mandibular component and fossa component in the implanted mandible. All the other contact pairs (implant-screw, mandible-screw, cortical-cancellous, teeth-PDL and PDL-cancellous) were defined as bonded. A coefficient of friction of 0.3 was chosen for contact interfaces since previous studies^35,36^ reported a negligible influence of coefficient of friction on the overall stress distribution for a properly fitted implanted model under physiological loading. An ‘Augmented Lagrangian Algorithm’ was defined for surface-to-surface contact pairs.

**Figure 3.**
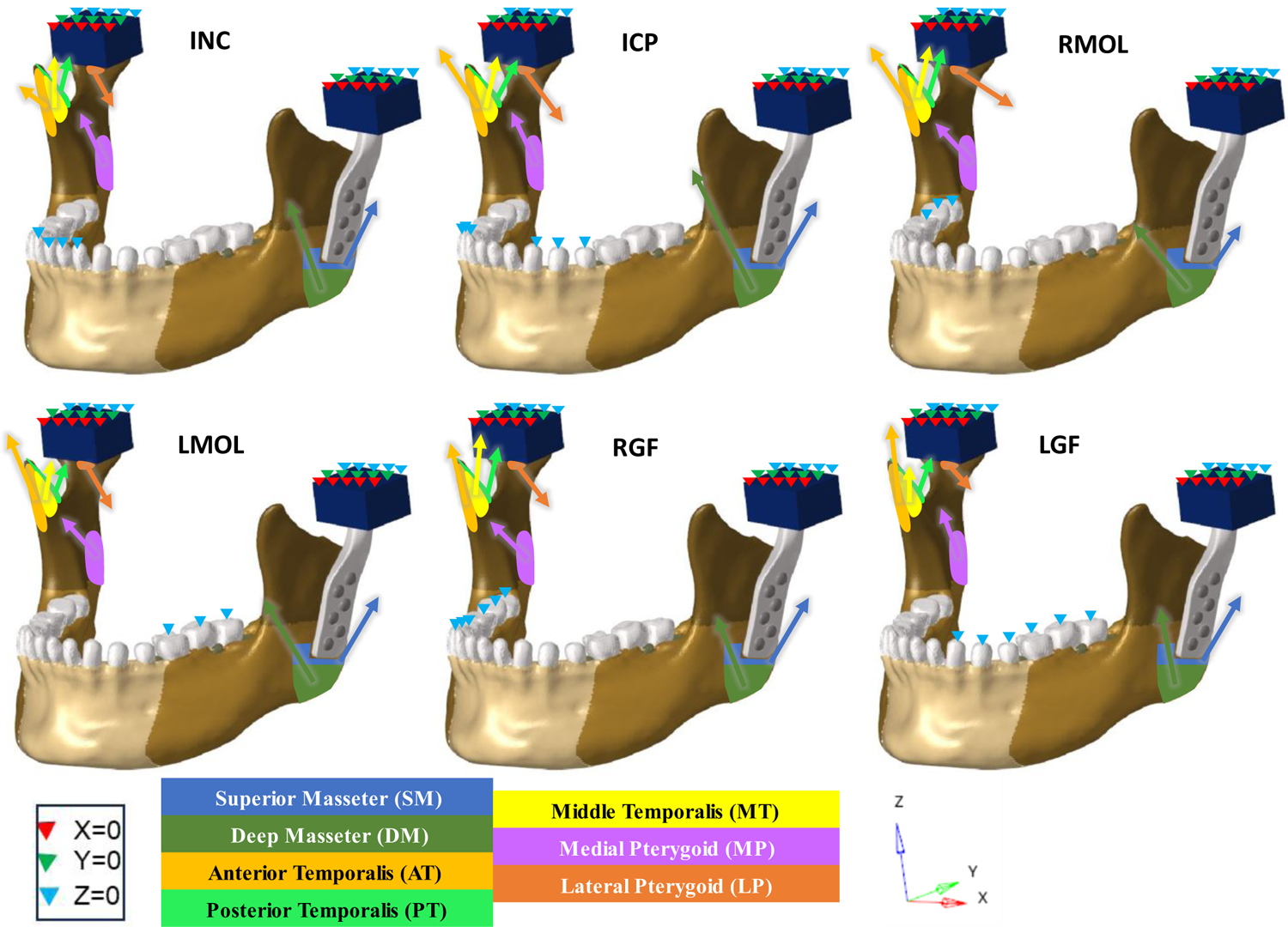
Muscle attachment sites, direction, and magnitude of force with boundary conditions on implanted mandible with narrow TMJ implant

**Table 2:** Muscle forces and boundary conditions simulating different clenching activities^19,34^.

| Clenching tasks | Side | Direction | Muscular force (N) |  |  |  |  |  |  | Boundary Condition |
| --- | --- | --- | --- | --- | --- | --- | --- | --- | --- | --- |
|  |  |  | SM | DM | MP | AT | MT | PT | LP |  |
| INC | Right | Force | 76.2 | 21.2 | 136.4 | 12.6 | 5.7 | 3.0 | 47.5 | Constrained the incisor regions |
|  |  | F<sub>x</sub> | -15.8 | -11.6 | 66.3 | -1.9 | -1.3 | -0.6 | 29.9 |  |
|  |  | F<sub>y</sub> | -31.9 | 7.6 | -50.9 | -0.6 | 2.9 | 2.6 | -36.0 |  |
|  |  | F<sub>z</sub> | 67.3 | 16.1 | 107.8 | 12.5 | 4.8 | 1.4 | -8.3 |  |
|  | Left | Force | 76.2 | 21.2 | 136.4 | 12.6 | 5.7 | 3.0 | 47.5 |  |
|  |  | F<sub>x</sub> | 15.8 | 11.6 | -66.3 | 1.9 | 1.3 | 0.6 | -29.9 |  |
|  |  | F<sub>y</sub> | -31.9 | 7.6 | -50.9 | -0.6 | 2.9 | 2.6 | -36.0 |  |
|  |  | F<sub>z</sub> | 67.3 | 16.1 | 107.8 | 12.5 | 4.8 | 1.4 | -8.3 |  |
| ICP | Right | Force | 190.4 | 81.6 | 132.9 | 154.9 | 91.8 | 71.0 | 18.1 | Constrained the canine and premolars |
|  |  | F<sub>x</sub> | -39.4 | -44.6 | 64.6 | -23.1 | -20.4 | -14.8 | 11.4 |  |
|  |  | F<sub>y</sub> | -79.80 | 29.2 | -49.61 | -6.8 | 45.9 | 60.8 | -13.7 |  |
|  |  | F<sub>z</sub> | 168.3 | 61.9 | 105.1 | 153.0 | 76.8 | 33.7 | -3.1 |  |
|  | Left | Force | 190.4 | 81.6 | 132.9 | 154.9 | 91.8 | 71.0 | 18.1 |  |
|  |  | F<sub>x</sub> | 39.4 | 44.6 | -64.6 | 23.1 | 20.4 | 14.8 | -11.4 |  |
|  |  | F<sub>y</sub> | -79.8 | 29.2 | -49.6 | -6.8 | 45.9 | 60.8 | -13.7 |  |
|  |  | F<sub>z</sub> | 168.3 | 61.9 | 105.1 | 153.0 | 76.8 | 33.7 | -3.1 |  |
| RMOL | Right | Force | 137.1 | 58.8 | 146.9 | 115.4 | 63.1 | 44.6 | 20.1 | Constrained the right molars |
|  |  | F<sub>x</sub> | -28.4 | -32.1 | 71.4 | -17.2 | -14.0 | -9.3 | 12.6 |  |
|  |  | F<sub>y</sub> | -57.4 | 21.0 | -54.8 | -5.1 | 31.5 | 38.1 | -15.2 |  |
|  |  | F<sub>z</sub> | 121.2 | 44.5 | 116.1 | 114.0 | 52.8 | 21.1 | -3.5 |  |
|  | Left | Force | 114.2 | 49.0 | 104.9 | 91.7 | 64.0 | 29.5 | 43.5 |  |
|  |  | F<sub>x</sub> | 23.6 | 26.7 | -51.0 | 13.7 | 14.2 | 6.1 | -27.4 |  |
|  |  | F<sub>y</sub> | -47.9 | 17.5 | -39.1 | -4.0 | 32.0 | 25.2 | -32.9 |  |
|  |  | F<sub>z</sub> | 101.0 | 37.1 | 83.0 | 90.5 | 53.6 | 14.0 | -7.6 |  |
| LMOL | Right | Force | 114.2 | 49.0 | 104.9 | 91.7 | 64.0 | 29.5 | 43.5 | Constrained the left molars |
|  |  | F<sub>x</sub> | -23.6 | -26.7 | 51.0 | -13.7 | -14.2 | -6.1 | 27.4 |  |
|  |  | F<sub>y</sub> | -47.9 | 17.5 | -39.1 | -4.0 | 32.0 | 25.2 | -32.9 |  |
|  |  | F<sub>z</sub> | 101.0 | 37.1 | 83.0 | 90.5 | 53.6 | 14.0 | -7.6 |  |
|  | Left | Force | 137.1 | 58.8 | 146.9 | 115.4 | 63.1 | 44.6 | 20.1 |  |
|  |  | F<sub>x</sub> | 28.4 | 32.1 | -71.4 | 17.2 | 14.0 | 9.3 | -12.6 |  |
|  |  | F<sub>y</sub> | -57.4 | 21.0 | -54.8 | -5.1 | 31.5 | 38.1 | -15.2 |  |
|  |  | F<sub>z</sub> | 121.2 | 44.5 | 116.1 | 114.0 | 52.8 | 21.1 | -3.5 |  |
| RGF | Right | Force | 34.3 | 29.4 | 12.2 | 104.3 | 61.2 | 46.8 | 39.5 | Constrained the right canine, premolars and molars |
|  |  | F<sub>x</sub> | -7.1 | -16.0 | 5.9 | -15.5 | -13.6 | -9.7 | 24.9 |  |
|  |  | F<sub>y</sub> | -21.5 | 7.6 | -49.6 | -0.5 | 2.9 | 3.9 | -7.1 |  |
|  |  | F<sub>z</sub> | 45.4 | 16.1 | 105.1 | 10.9 | 4.8 | 2.2 | -1.6 |  |
|  | Left | Force | 51.4 | 21.2 | 132.9 | 11.1 | 5.7 | 4.5 | 9.4 |  |
|  |  | F<sub>x</sub> | 10.6 | 11.6 | -64.6 | 1.6 | 1.3 | 0.9 | -5.9 |  |
|  |  | F<sub>y</sub> | -21.5 | 7.6 | -49.6 | -0.5 | 2.9 | 3.9 | -7.1 |  |
|  |  | F<sub>z</sub> | 45.4 | 16.1 | 105.1 | 10.9 | 4.8 | 2.2 | -1.6 |  |
| LGF | Right | Force | 51.4 | 21.2 | 132.9 | 11.1 | 5.7 | 4.5 | 9.4 | Constrained the left canine, premolars and molars |
|  |  | F<sub>x</sub> | -10.6 | -11.6 | 64.6 | -1.6 | -1.3 | -0.9 | 5.9 |  |
|  |  | F<sub>y</sub> | -21.5 | 7.6 | -49.6 | -0.5 | 2.9 | 3.9 | -7.1 |  |
|  |  | F<sub>z</sub> | 45.4 | 16.1 | 105.1 | 10.9 | 4.8 | 2.2 | -1.6 |  |
|  | Left | Force | 34.3 | 29.4 | 12.2 | 104.3 | 61.2 | 46.8 | 39.5 |  |
|  |  | F<sub>x</sub> | 7.1 | 16.0 | -5.9 | 15.5 | 13.6 | 9.7 | -24.9 |  |
|  |  | F<sub>y</sub> | -21.5 | 7.6 | -49.6 | -0.5 | 2.9 | 3.9 | -7.1 |  |
|  |  | F<sub>z</sub> | 45.4 | 16.1 | 105.1 | 10.9 | 4.8 | 2.2 | -1.6 |  |

### 2.5 Interpretation of Results

The decision to primarily analyse the cortical bone stems from its critical role in load transfer during mastication. Cortical bone bears the majority of the load compared to trabecular bone^37^. The primary focus of this study was to understand the failure in the mandible and implant. Given that the mandible is a sandwich-like structure encapsulating softer cancellous bone, its failure is primarily governed by the failure of cortical bone. Cortical bone being brittle in nature, prior studies considered maximum principal strains as failure criterion ^38,39^. Therefore, in the current study, maximum principal strains and stresses in the mandible were considered for biomechanical comparisons between the intact and implanted mandibles. To exclude singular values, owing to constrained regions in the FE models in both intact and implanted mandibles, 99^th^ percentile values of principal stresses and strains of cortical bone were considered for comparison. Conversely, implants being made of Ti6Al4V which exhibits ductile material behaviour, von Mises stress was considered for investigating the load transfer through narrow and standard TMJ implants under unilateral and bilateral clenching.

### 2.6 Verification of Intact FE model

Since the CT-scan dataset was derived from a living subject, the FE model developed in the present study could not be directly validated by a one-to-one experiment. However, a qualitative and quantitative verification of the FE model was performed by comparing the results with those from earlier studies ^12,16,24,40^. The present study observed a maximum principal stress of 11.87 MPa around the biting surfaces of the right molar teeth which corroborates a similar finding (15 MPa to 25 MPa) reported in a previous study by Korioth et al. ^12^ during a right molar clenching. The present study observed high maximum principal strains [700-1750 με] in all the load cases in healthy mandible. The magnitude of the principal strain observed near the anterior ramus surface of the mandible during the RMOL case in the present study was 1250 με, which closely matches the strain (1260 με) observed earlier ^24^. Similar to the study by ^40^ high tensile strains below the mandibular notch and anterior surface of the condylar neck were also observed here (Fig 3). Gröning et al. ^16^ reported highest principal strains around the *pars alveolaris* below the molars, oblique line, *basis mandibulae,* and posterior ramus inferior to condyle during INC, ICP and molar bites (RMOL/LMOL). Similar strain distributions were observed in the present study for INC, ICP, and molar bites. The quantitative difference in results in maximum principal stresses and strains between this study and the erstwhile works ^12,24^ might be attributed to the difference in bone morphology, choice of bone material properties, and in muscle attachment sites. Notwithstanding subtle differences, the results from the present study provide confidence in the developed FE models.

## 3. Results

The principal stress and strain on the intact and implanted mandibles along with von Mises stress in the TMJ implants (mandibular component and screws) during immediate post-operative non-osseointegrated (NOI) condition are discussed first (section 3.1 and 3.2). Thereafter, influence of osseointegration on stresses and strains has been investigated by comparing the non-osseointegrated results with those of the osseointegrated (OI) models (section 3.3).

### 3.1 Maximum Principal Stress and Strain Distribution in Intact and Implanted Mandibles

Figure 4 and 7a shows the magnitude and distribution of maximum principal stress around intact and implanted mandibles under six different biting tasks respectively. The maximum tensile stress (Fig. 7a) in the intact mandible ranges between 6 MPa (LGF) and 16 MPa (ICP). Higher stress regions were observed along the oblique line, anterior ramus, and across lateral alveolar regions in discrete patches for the molar (RMOL) and intercuspal (ICP) occlusion. For the rest of the load cases (INC/LGF/RGF), higher stresses were observed mostly near the molars in the alveolar region and around the medial mandibular notch (Fig. 4). Minor difference in stress magnitude (narrow TMJ: 1.02 MPa; standard TMJ: 0.75 MPa) was observed between the intact and the implanted mandibles. However, marginally higher stress (maximum difference of 0.32 MPa) was observed in mandible with standard TMJ implant as compared to the one with narrow TMJ implant. Moreover, an overall increase in stress was observed in case of contralateral clenching (RMOL and RGF) as compared to ipsilateral ones (LMOL and LGF).

**Figure 4.**
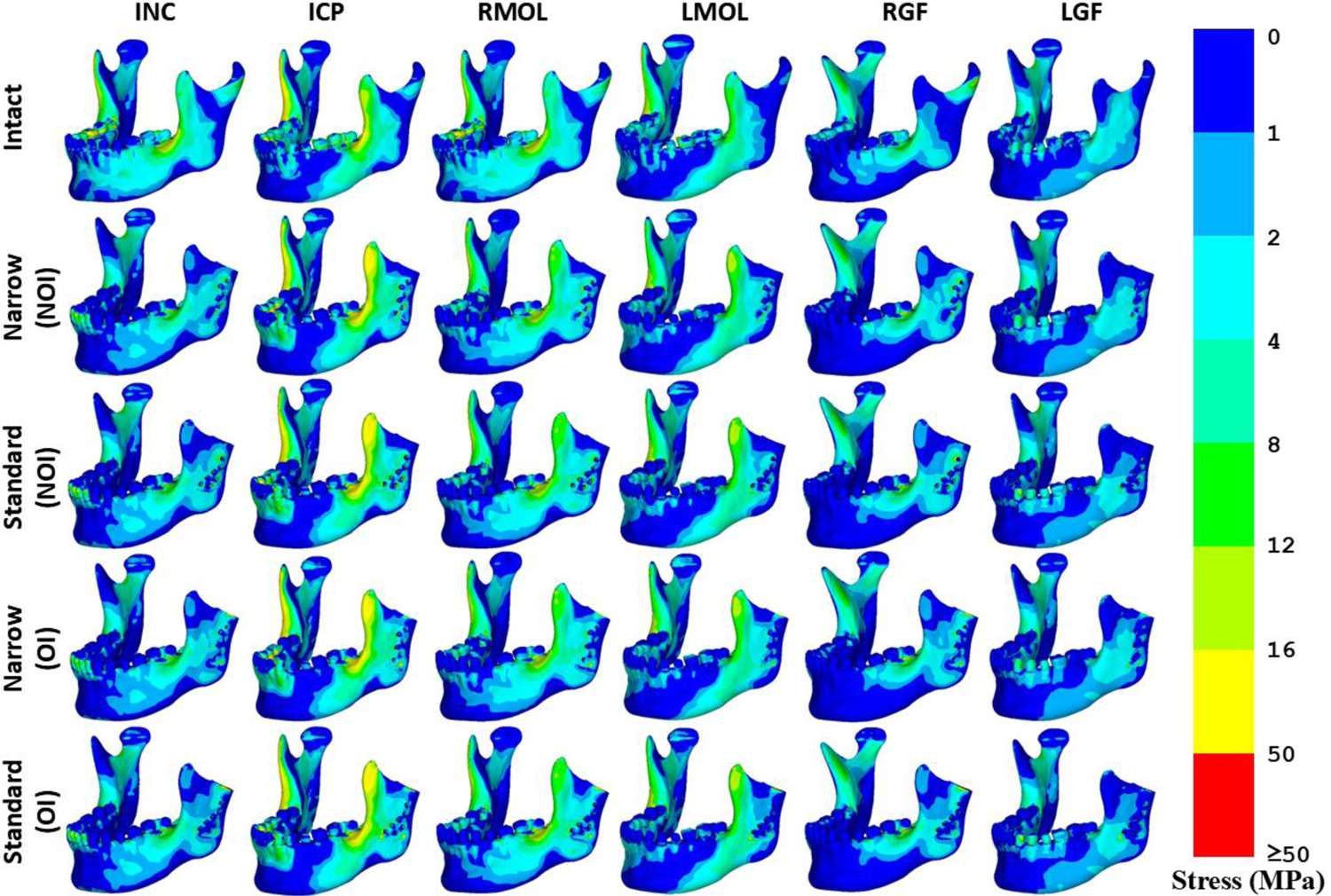
Maximum principal stress distributions across intact and implanted mandibles under unilateral and bilateral clenching. For implanted mandibles, stress distributions for both non-osseointegrated (NOI) and osseointegrated (OI) conditions are shown.

Figure 5 and 7b shows the distribution and magnitude of maximum principal strain in the intact and reconstructed mandibles under unilateral and bilateral clenching respectively. The strain in the intact mandible over all the load cases ranges from 700 με to 1750 με (Fig. 7b). Higher strains (1150 με to 1750 με) were observed during molar biting (RMOL and LMOL) and ICP cases across the ramus primarily around the lower condyles, coronoid process, oblique line and lateral surfaces of alveolar and body (Fig. 5). However, INC and group function tasks (RGF/LGF) resulted in lower strains (700 με to 730 με) mostly around medial surfaces of mandibular notch and alveolar region on lateral sides. The ICP clenching caused the highest tensile strain (1750 με to 1880 με) among all the load cases.

**Figure 5.**
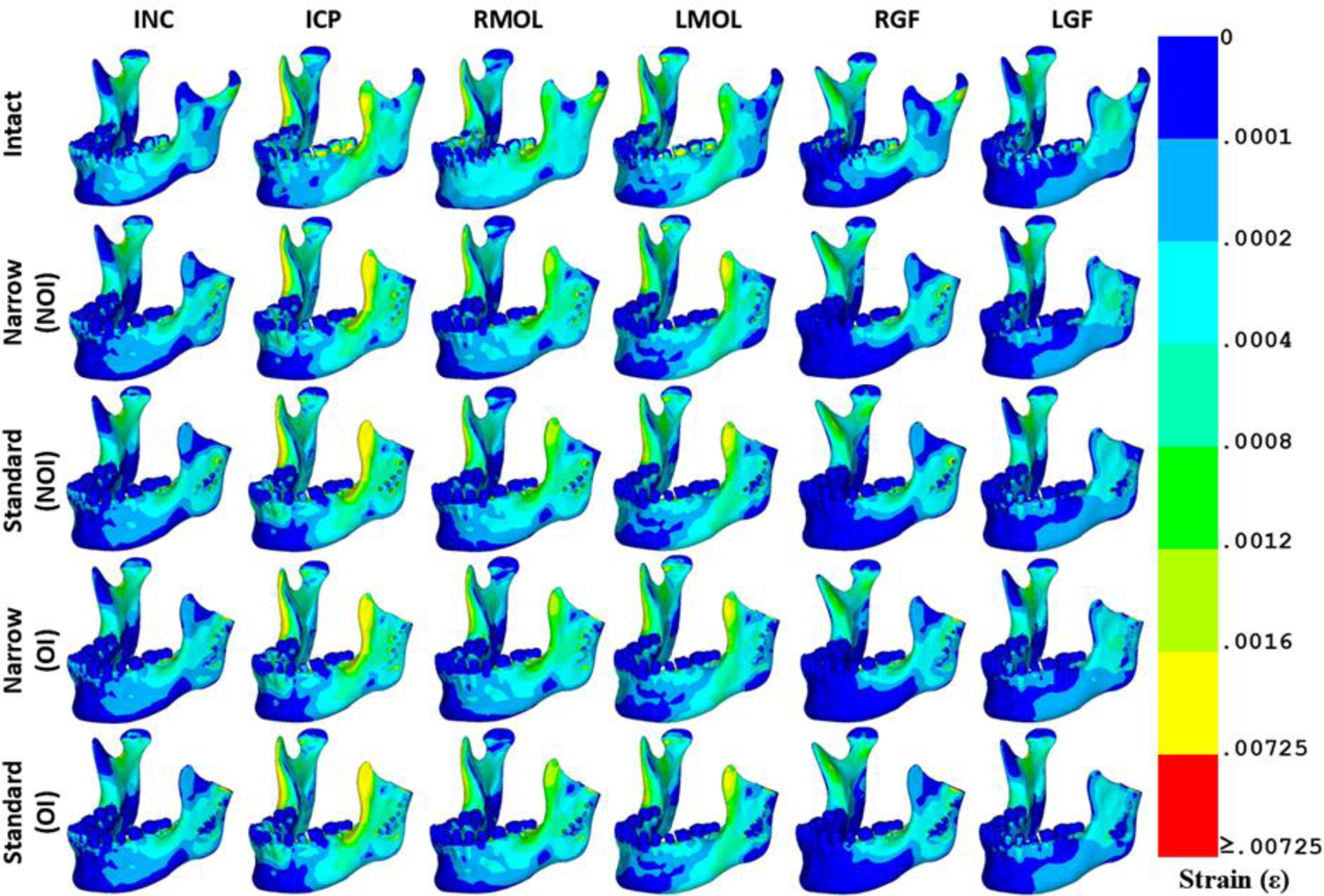
Maximum principal strain distribution across the intact and implanted mandibles under unilateral and bilateral clenching. For implanted mandible, strain distributions during both non-osseointegrated (NOI) and osseointegrated (OI) conditions are shown.

An overall reduction in the magnitude of peak strain was observed in implanted mandibles (narrow TMJ: 80 με; standard TMJ: 120 με) as compared to those in intact mandible. However, higher strains (up to 80%) were observed around the screw holes in the implanted ones as compared to the intact mandible. Unlike intact mandibles, maximum principal strains in implanted mandibles increased under ipsilateral clenching (up to 14.75%) than contralateral ones (Figure 7b). However, an opposite trend was observed for ipsilateral gonial angle region wherein strains under ipsilateral loading was higher (by 11.9%) than those under contralateral loading.

### 3.2 Stresses in Implants

Figure 6 and 8 shows the von Mises stress distribution and peak von Mises stress across the implants and screws under different biting conditions respectively. Among all the loading conditions, peak von Mises stress in the mandibular component (Fig. 8a) of ramus standard TMJ was the highest (151 MPa) under RMOL, whereas the lowest (58 MPa) under LMOL. In comparison, for narrow TMJ implant, the highest (132 MPa) and lowest (87 MPa) von Mises stresses in mandibular component were observed under RGF and LGF, respectively. In general, ipsilateral clenching (LMOL/LGF) resulted in reduced von Mises stresses (standard TMJ: 93 MPa; narrow TMJ: 45 MPa) in mandibular component as compared to contralateral clenching (RMOL/RGF). Regions near the base of the mandibular component experienced negligible stresses (≤5MPa in standard TMJ implants (Fig. 6). Peak von Mises stresses in the screws (Fig. 8b) were minimum during LMOL (narrow TMJ: 50 MPa; standard TMJ: 52 MPa). In comparison, higher stresses in the screws were observed during INC (narrow TMJ: 115 MPa; standard TMJ: 128 MPa) and RGF (narrow TMJ: 109 MPa; standard TMJ: 130 MPa). Additionally, stresses in the screws were higher in standard TMJ implant as compared to narrow TMJ implant during immediate post-operative non-osseointegrated condition. Overall stress patterns in the screws under all the loading conditions were similar.

**Figure 6.**
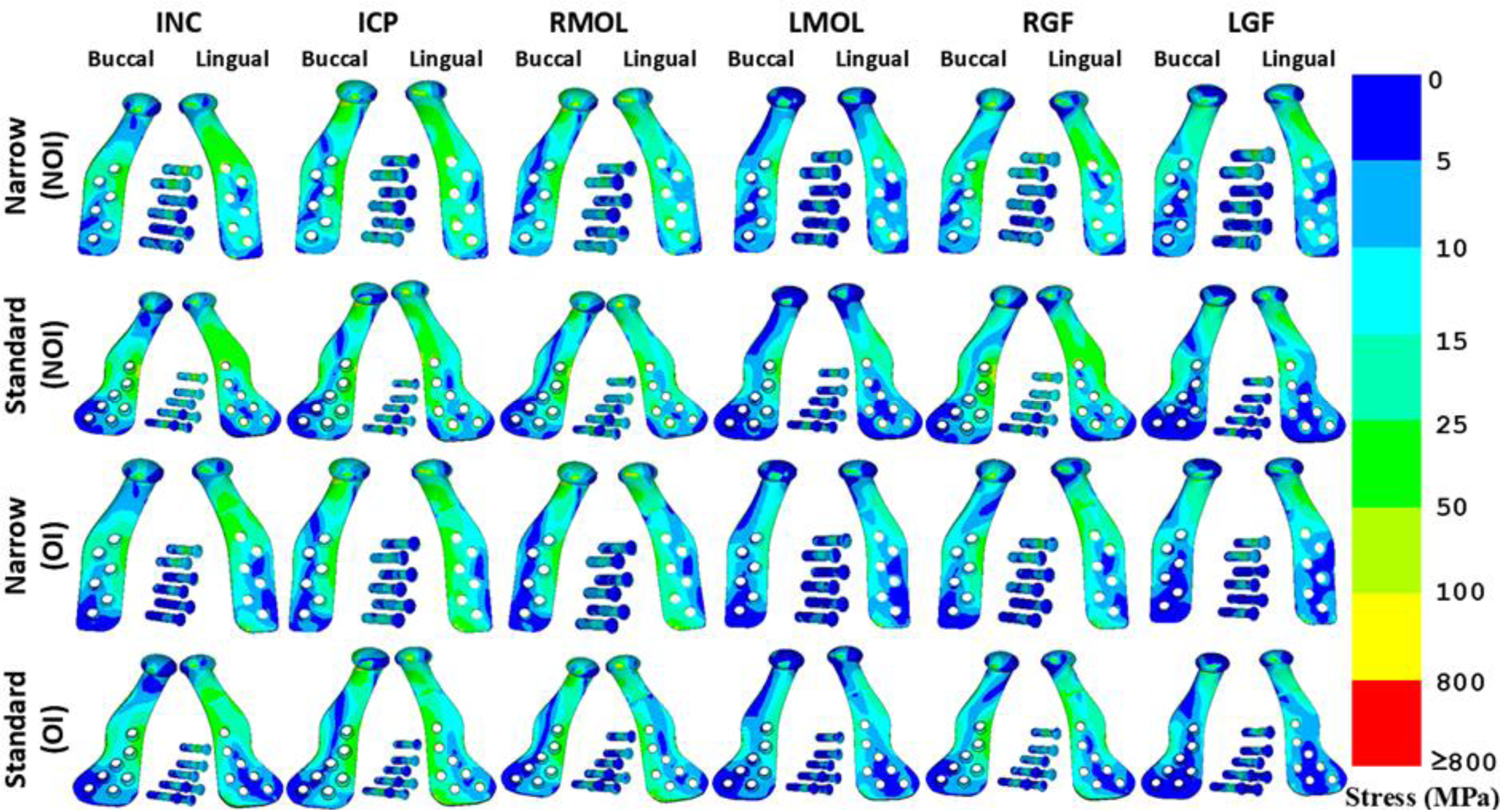
von Mises stress distribution in narrow and standard TMJ implants and screws under unilateral and bilateral clenching during non-osseointegrated (NOI) and osseointegrated (OI) conditions.

**Figure 7.**
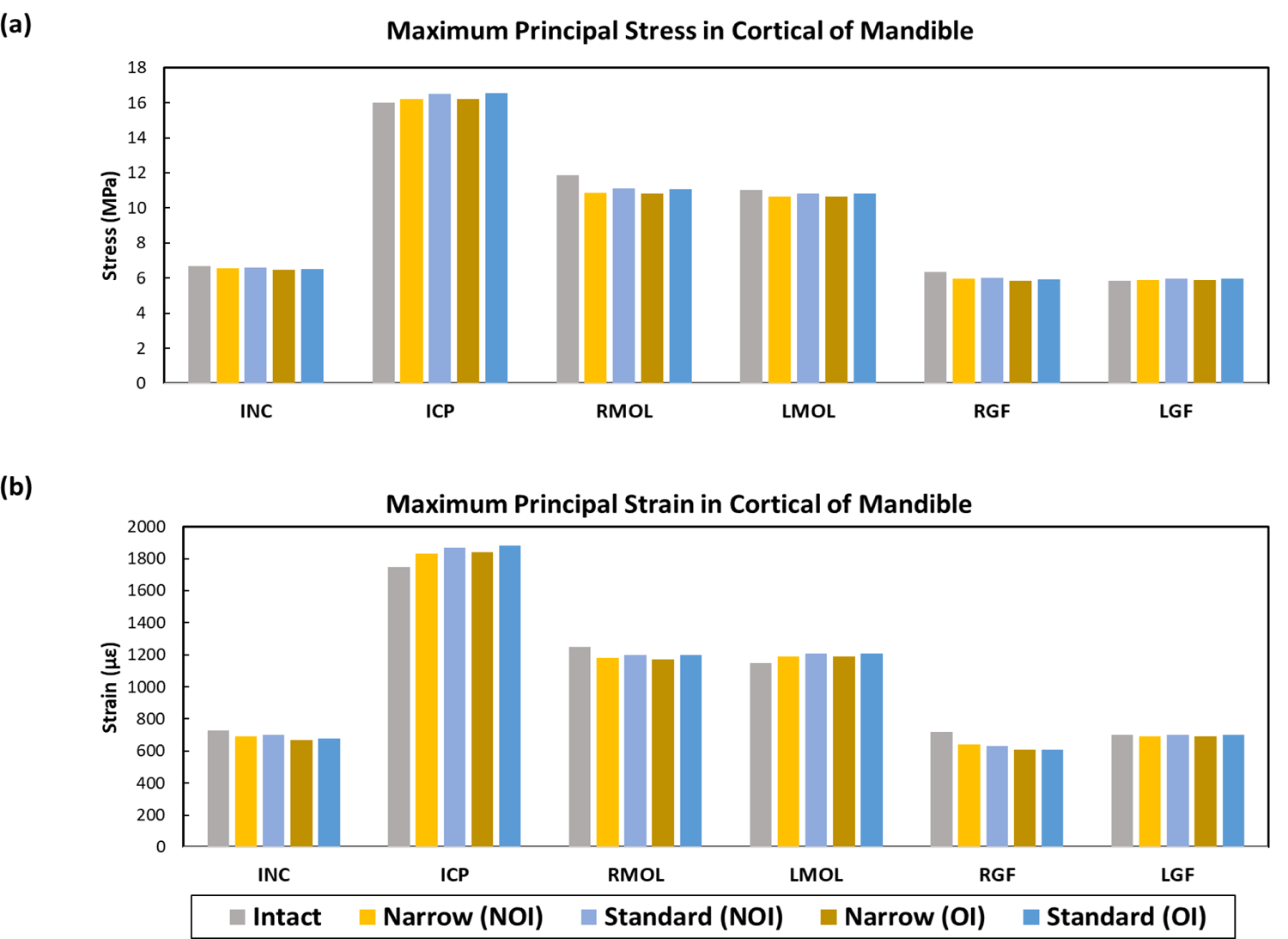
(a) Maximum principal stress, and (b) maximum principal strain in the cortical bone of intact and implanted (with narrow/standard mandibular component) mandible.

**Figure 8.**
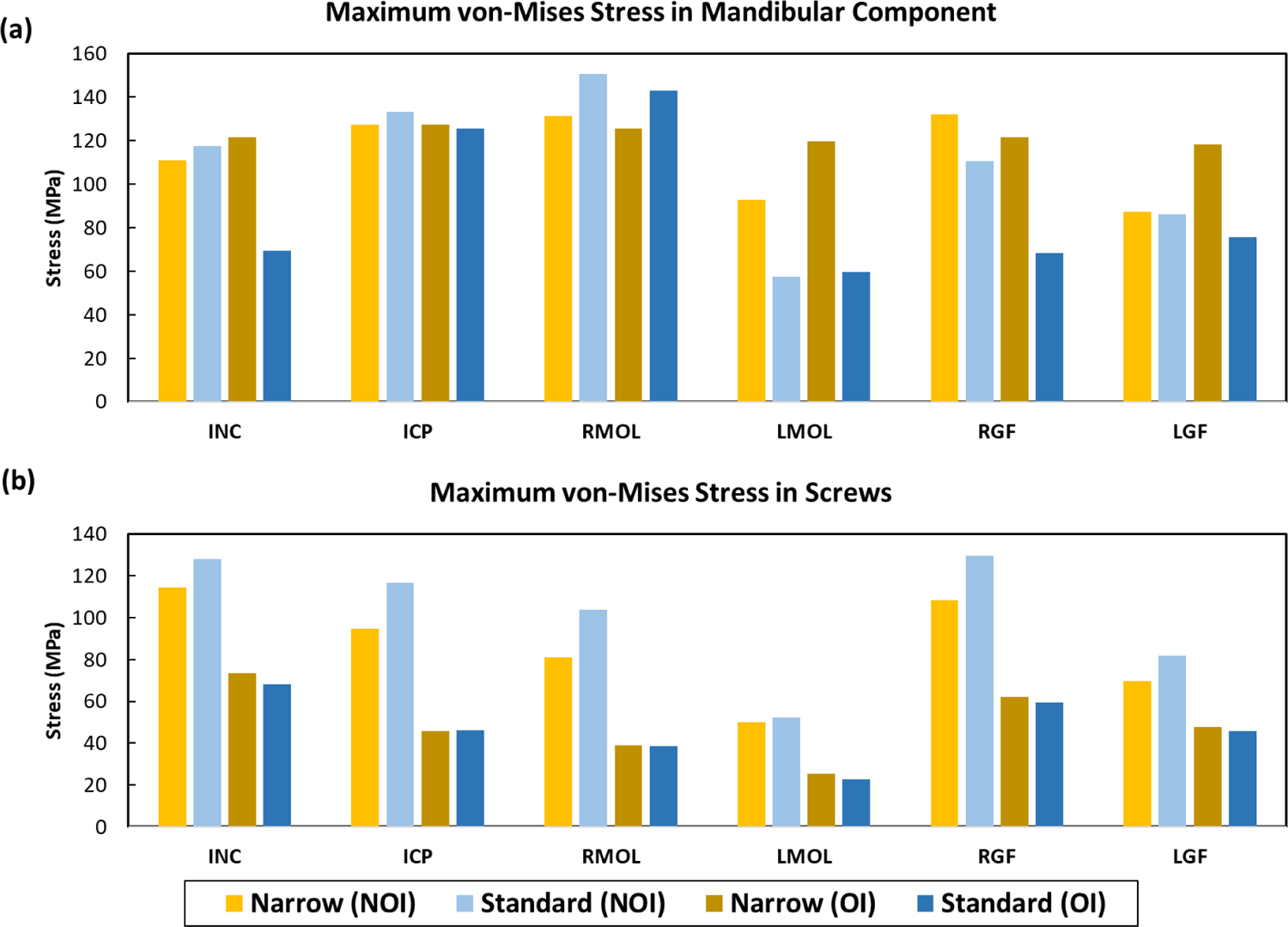
Maximum von Mises stresses in (a) mandibular component and (b) screws.

### 3.3 Influence of osseointegration on stresses and strains in implanted mandible

The present study investigated both immediate post-operative non-osseointegrated (NOI) and long-term post-operative osseointegrated (OI) conditions. The interfacial conditions were found to influence the stresses (Figure 4 and 7a) and strains (Figure 5 and 7b) in the mandible and implant. A marginal reduction in mandibular stresses (narrow TMJ: 0.14 MPa; standard TMJ: 0.12 MPa) and strains (narrow TMJ: 30 µε; standard TMJ: 20 µε) were observed due to osseointegration. However, in mandibular component, an increase in stress was observed under contralateral loading as compared to ipsilateral loading for both NOI and OI conditions. During NOI condition, von Mises stresses in mandibular component during LMOL (narrow TMJ: 93 MPa; standard TMJ: 58 MPa) and LGF (narrow TMJ: 87 MPa; standard TMJ: 86 MPa) were found to be much lower than those during RMOL (narrow TMJ: 131 MPa; standard TMJ: 151 MPa) and RGF (narrow TMJ: 132 MPa; standard TMJ: 111 MPa) loading conditions. A similar observation was noted for OI condition as well (Figure 8). Osseointegration also reduced stresses in mandibular components (up to 48 MPa) and screws (up to 71 MPa). Notably, all observed stress values remained well below the yield limit of the implant material (∼955 MPa)^41^ (Fig. 7a).

## 4. Discussion

The present study is aimed to assess biomechanical performance of two widely used stock mandibular TMJ implants (standard and narrow) under unilateral and bilateral clenching activities. In addition, influences of osseointegration on the stresses and strains in mandible and TMJ implants have also been investigated.

Minor difference in stress magnitude across the mandible between the intact and implanted mandibles was observed in this study. Although mandibles implanted with standard TMJ implants resulted in slightly higher stresses (0.05-0.34 MPa) in cortical bone, the overall spatial distribution of cortical bone stresses were similar in all the cases. Higher cortical bone strains were, however, observed around the screw holes. This might cause fibrous tissue formation at the bone-implant interface ^42, 43^ in the long term. Similar observation was also reported in earlier clinical study ^44^. Sugiura et al.^45^ reported a threshold compressive strain (3600 με) for inducing bone resorption in human mandibles. The peak strain reported in the present study in implanted mandibles with both implants is much below the critical threshold limit for bone resorption which ensures the suitability of these implants in the present scenario. Higher strains (up to 14.75%) on implanted bone were observed under ipsilateral loading as compared to the contralateral ones, which is in contrast with the observations for intact bone. A similar trend of strains under ipsilateral and contralateral loading was also reported earlier ^24^.

Bone-implant interface condition was found to influence load transfer through mandibular component. An overall reduction in stresses was observed due to osseointegration, similar to a recent study on dental implants ^29^. During the initial NOI condition, screws act as mechanism to transfer loads from the mandible to the mandibular component and vice versa. On the contrary, during the OI condition, the entire bone-implant interface becomes osseointegrated and therefore, load-sharing happens through a larger interfacial area. Hence, a reduction in peak von Mises stress in implant and screws due to osseointegration is observed. However, the marginal differences in mandibular stresses and strains between OI and NOI conditions can be attributed to the rigid fixation of the TMJ implant using bicortical screws, which limits implant-bone micromotion even in the immediate postoperative period. Furthermore, the implant’s rough surface promotes osseointegration, leading to additional stabilisation over time. Given these factors, the assumed coefficient of friction (µ = 0.3) at the implant-bone interface resulted in micromotion levels below the threshold (<5.4 micron) that could compromise osseointegration. Consequently, the frictional and bonded conditions exhibited negligible influence on the overall mandibular stress and strain distribution, as also supported by a previous study^46^. Although implant and screw stresses remained within the corresponding yield limit during both non-osseointegrated and osseointegrated (OI) conditions, during non-osseointegrated condition, screw stresses were higher in standard TMJ implant than those of the narrow TMJ implant. Therefore, based on this study, although both the stock implants were found to be safe, narrow TMJ is preferred over standard TMJ for joint replacement.

There were certain limitations of the present study. Even though the bone is heterogeneous anisotropic, the material property of cortical and cancellous bone was considered as a region-specific orthotropic material in the current work^12^. Modelling screws as cylindrical entities, while neglecting its threads, is another limitation of the present study. However, detailed thread modelling is primarily necessary when the focus is on understanding the stress-strain distribution at the screw thread interface ^47,48^ and, therefore, this assumption will not influence the overall prediction of this current study. In addition, inter-patient anatomical and muscle force variations of the human mandible were neglected in this study. Despite these limitations, this study on stock TMJ implants provide a thorough understanding on the performance of different stock implants under clenching loads and is expected to assist clinicians to choose a proper TMJ implant from biomechanical perspectives.

## 5. Conclusions

To summarize, the present study investigated the biomechanics of narrow and standard TMJ implants on a patient-specific human mandible under unilateral and bilateral biting tasks. Ipsilateral clenching resulted in higher strains in implant as compared to contralateral clenching. Overall, stresses and strains in the cortical bone, TMJ implants and screws were well below the corresponding failure limits for all the models. While osseointegration reduced stresses and strains in cortical bone for both the stock implants, a standard TMJ implant induced higher stress in cortical bone and higher strains in screws as compared to narrow TMJ implant during immediate post-operative non-osseointegrated condition. Therefore, this study suggests that although both standard and narrow TMJ implants were found to be safe for joint replacements, narrow TMJ implant should perhaps be preferred over the standard ones.

## Statements of ethical approval

Ethical approval was obtained from the Institute Ethics Committee (IEC) of All India Institute of Medical Sciences, New Delhi, India with the following approval numbers: IEC-611/15.07.2022, AA-3/05.08.2022, RP-11/2022.

## Acknowledgements

The authors would like to acknowledge the Department of Mechanical Engineering, IIT Delhi for providing the required computational facilities and AIIMS, New Delhi for providing the implants to carry out this research work.

## CRediT authorship contribution statement

**Rajdeep Ghosh:** Conceptualization, Data Curation, Formal Analysis, Investigation, Methodology, Software, Validation, Visualization, Writing – original draft, Writing – review & editing. **Girish Chandra:** Conceptualization, Data Curation, Formal Analysis, Methodology, Visualization, Writing – review & editing. **Vivek Verma:** Data Curation, Visualization, Writing – review & editing. **Kamalpreet Kaur:** Writing – review & editing. **Ajoy Roychoudhury:** Funding acquisition, Project Administration, Resources, Supervision. **Sudipto Mukherjee:** Conceptualization, Funding acquisition, Methodology, Project Administration, Resources, Supervision, Writing – review & editing. **Anoop Chawla:** Conceptualization, Funding acquisition, Methodology, Project Administration, Resources, Supervision, Writing – review & editing. **Kaushik Mukherjee:** Conceptualization, Funding acquisition, Methodology, Project Administration, Resources, Supervision, Writing – review & editing.

